# Ergothioneine, a metabolite of the gut bacterium *Lactobacillus reuteri*, protects against stress-induced sleep disturbances

**DOI:** 10.1101/2020.03.26.009423

**Authors:** Yoshiki Matsuda, Nobuyuki Ozawa, Takiko Shinozaki, Ken-ichi Wakabayashi, Kosuke Suzuki, Yusuke Kawano, Iwao Ohtsu, Yoshitaka Tatebayashi

## Abstract

The relationships between depression and gut microbiota, particularly those involving the immune system, have become a major focus of recent research. Here, we analyzed changes in gut microbiota and their sulfur metabolites in the feces of a depression rat model using the modified 14-day social defeat stress (SDS) paradigm. Our results showed that SDS increased fecal *Lactobacillus reuteri* in correlation with ergothioneine levels at around day 11, which continued for at least one month following SDS administration. In vitro study further revealed that *L. reuteri* is capable of producing ergothioneine. Although the known anti-inflammatory and anti-oxidative actions of ergothioneine suggested that the increased fecal ergothioneine levels may be related to intestinal anti-inflammatory defense mechanisms, no change was observed in the plasma ergothioneine levels during the same observation period, indicating that the defense mechanisms may not be sufficiently reflected in the body. As ergothioneine is a natural ingredient that is absorbed mainly from the upper gastrointestinal tract, we hypothesized that oral ergothioneine may exert antidepressant effects. As expected, oral administration of ergothioneine prior to and during the SDS paradigm had a preventative effect on SDS-induced depressive behaviors, such as social avoidance and depression-like sleep abnormalities, particularly those of rapid eye movement sleep. These findings indicate that ergothioneine, a metabolite of *L. reuteri*, may be a common substance in the microbiota-gut-brain axis that prevents stress-induced sleep disturbances, especially those associated with depression.

## Introduction

Psychosocial stress is an environmental factor associated with the increased incidence of psychiatric illnesses such as major depressive disorder (MDD) [1]. In addition to the core symptoms (depressed mood and loss of interest/pleasure), MDD can also be characterized by somatic symptoms such as sleep abnormalities [2]. We have recently developed a depression rat model using the 14day social defeat stress (SDS) paradigm [3]. Compared to other rodent stress models, nearly all SDS rats exhibit long-term social avoidance and MDD-like sleep abnormalities in our paradigm, which can be rescued by chronic antidepressant treatments [3]. In particular, the observed sleep abnormalities exhibit numerous similarities to those in patients with MDD. These include significantly increased rapid eye movement (REM) and decreased non-REM (NREM) sleep durations, increased sleep fragmentation, and decreased REM sleep latency during the light phase [3].

Recent research has reported the influence of gut microbiota on cerebral function (i.e., the microbiota-gut-brain axis) [4–9]. For example, depressive behaviors develop in germ-free rodents following fecal transplants from human patients with MDD [10, 11]. On the other hand, decreases in MDD-like behaviors occurred following the administration of prebiotics in mice subjected to mild chronic stress [12]. More recently, Pearson-Leary et al. demonstrated that the gut microbiota regulate the increases in MDD-like behaviors and inflammatory processes in the ventral hippocampus of SDS-vulnerable rats [13]. Nevertheless, considerably less research is available regarding stress-induced changes in gut bacteria-produced metabolites, especially sulfur metabolites, or their effects on cerebral functions.

Despite its essential involvement in all organisms, our understanding of sulfur metabolism lags behind that of carbon or nitrogen, mostly due to difficulty in metabolite detection. Cellular sulfur metabolites are quantitatively scarce and readily undergo redox reactions on their thiol group. Recently, however, the combination of sensitive liquid chromatography-tandem mass spectrometry (LC-MS/MS) with thiol-specific derivatization methods using monobromobimane has enabled such detection [14–17]. Here, we analyzed how SDS influences gut microbiota and their sulfur metabolites in rats and found that fecal *Lactobacillus reuteri* increased concomitantly with ergothioneine during the late SDS stage. To evaluate the role of ergothioneine, we preventatively administered ergothioneine to the SDS rats prior to and during SDS and analyzed its effects on the depressive behaviors and sleep abnormalities in our SDS rat model.

## Materials and Methods

### Animals

All procedures were approved by the Animal Use and Care Committee of the Tokyo Metropolitan Institute of Medical Science for Ethics of Animal Experimentation. Animals were kept under standard laboratory conditions [12 h light/dark cycle, lights on at 08:00 (= Zeitgeber time 0; ZT0)] with food and water available *ad libitum* unless otherwise indicated. All animal experiments were performed in 2015-2018.

### Social defeat paradigm

We applied a modified repeated SDS model [3] for 14 consecutive days using male Sprague Dawley (SD) rats that were approximately 8 weeks old (Charles River Laboratories Japan, Yokohama, Japan) at stress onset. Briefly, each SD rat was transferred into the home cage of a retired aggressive male Brown Norway (BN) rat (>7 months of age; Charles River Laboratories Japan). The resident BN rat was allowed direct physical contact with the SD rat (intruder) for 10 min, and then resident and intruder rats were kept in indirect contact for 24 h in a resident cage using a perforated clear divider to prevent physical contact. The next day, the intruder was exposed to a novel resident BN aggressor. Subsequently, intruders were subjected to combined stress (direct and indirect contact) for the first five weekdays, followed by only indirect contact for the subsequent two weekend days. This process was continuously repeated for two weeks. Control animals were housed on one side of a perforated divider without a resident. Upon termination of SDS, all rats were housed individually.

### Social interaction

The social interaction test was conducted to assess social avoidance behavior [3, 18, 19]. Briefly, the arena was an open field (90 × 90 × 45 cm) maintained in weak lighting conditions (30 lux). An experimental SD rat was placed inside the arena, and its movements were monitored using an infrared camera for two consecutive sessions of 2.5 min each. During the first session (“No target”), an empty wire mesh cage (30 × 15 × 15 cm) was placed at one end of the field. During the second session (“target”), an unfamiliar BN or SD rat was placed in the mesh cage. Time spent in the interaction zone was quantified using custom applications (Time OFCR4, O’Hara & Co., Tokyo, Japan). The interaction ratio was calculated as (interaction time, “target”) / (interaction time, “No target”) × 100 %.

### Fecal and blood sample collection

Fresh fecal samples were collected before the first SDS application (before), 1 day (stress 2d), 4 days (stress 5d), and 10 days (stress 11d) after the first SDS application, and 1 day (after stress), 7 days (1W), and 1 month (1M) after the last SDS, and stored at −80 °C until further analysis. After the end of SDS periods, blood plasma samples were collected via cardiac puncture under pentobarbital sodium (Somnopentyl; Kyoritsu, Tokyo, Japan) anesthesia (60 mg/kg, intramuscular) for subsequent metabolite analysis.

### 16S rRNA gene sequencing analysis

Bacterial genomic DNA in fecal samples was extracted, and then two-step polymerase chain reactions (PCRs) were performed. The first PCR utilized either universal primer pair corresponding to the V3–V4 region of the 16S rRNA gene: (1) 341F (5’-CCTACGGGNGGCWGCAG-3’) and 805R (5’-GACTACHVGGGTATCTAATCC-3’) or (2) 341F (5’-TCGTCGGCAGCGTCAGATGTGTATAAGAGACAGCCTACGGGNGGCWGCAG-3’) and 806R (5’-GTCTCGTGGGCTCGGAGATGTGTATAAGAGACAGGGACTACHVGGGTWTCTAAT-3’) (TaKaRa Bio, Shiga, Japan). The second PCR was performed to add the index sequences for Illumina sequencing with barcode sequences. The prepared libraries were subjected to either paired-end 300 [for (1)] or 250 [for (2)] base sequencing, using the MiSeq Reagent Kit v3 on the MiSeq (Illumina, San Diego, CA, USA).

Microbiome analysis was performed using the microbial genomics module either on the CLC Genomics Workbench version 8.5.1 (Qiagen, Hilden, Germany) [for (1)] or the open-source bioinformatics pipeline Quantitative Insights Into Microbial Ecology (QIIME) version 1.8.0 [for (2)]. Sequenced paired-end reads were assembled to construct contigs; chimeric contigs were removed by applying either the chimera crossover detection algorithm [for (1)] or CD-HIT-operational taxonomic unit (OTU) algorithm [for (2)]. The remaining contigs were clustered into OTUs with 97% sequence similarity. To acquire taxonomic information for each OTU, representative sequences were assigned to the Greengenes 16S rRNA database by RDP classifier (version 2.2).

### Analysis of sulfur metabolomics

Sulfur metabolomics (Sulfur index) was performed using S-bimanyl derivatives via LC-MS/MS (Shimadzu Nexera UHPLC system with on-line LCMS 8040, Kyoto, Japan) as described previously [13–16]. Briefly, the sulfur-containing compounds were extracted from samples by adding methanol and converted to fluorescent derivatives using a thiol-specific alkylating reagent (mono-bromobimane). The target metabolite levels were determined from the peak area by mass chromatography and represented as relative amounts following normalization using the internal standard (D-camphor-10-sulfonic acid) peak area.

### In vitro study

*L. reuteri* (JCM Nos. 1112, 5867,1081, 1084, 5868, 5869, RIKEN BRC, Japan) and *Escherichia coli* (DH5α, BW2) were cultured in GAM medium at 30 °C for 24 h. Bacteria and supernatants were then collected for Sulfur index analysis.

### Oral ingestion of L-ergothioneine

Oral administration of L-ergothioneine (0.25 mg/ml; Nagase & Co., Tokyo, Japan) aqueous solution was conducted from 1 week prior to SDS initiation (day −7) to the end SDS application (day 14). The concentration of L-ergothioneine administered corresponds to approximately 30 mg/kg/day based on the initial water intake and body weight. Tang et al. demonstrated that daily oral administration of ergothioneine (35 or 70 mg/kg/day for 1, 7, and 28 days) induces accumulation of ergothioneine in whole-body tissues, including the brain, in male C57BL6J mice [20]. Control animals received only water.

### Surgery and electroencephalogram (EEG) recording

Under pentobarbital sodium anesthesia (60 mg/kg, intramuscular), the rats were fixed to a stereotaxic apparatus (SR-6M; Narishige, Tokyo, Japan). Non-polarized Ag/AgCl screw electrodes (1 mm diameter, O’Hara & Co., Tokyo, Japan) were implanted epidurally on the left side of the parietal cortex (LPa; Br −2.0, L1.5). Reference and ground electrodes were placed above the cerebellum. An electromyography (EMG) stainless electrode surface was subcutaneously placed on the dorsal neck muscle. The lead wires from all electrodes were soldered to a small socket and mounted on the skull with acrylic resin cement, along with these electrodes. A recovery period (> 7 days) was scheduled before initiation of the experiment.

To collect the EEG data, the rat was moved to an experimental cage with a soundproof box in which the 12 h light/dark cycle was maintained and connected via a recording cable with a built-in operational amplifier (TL074; Texas Instruments, Dallas, TX, USA) to reduce electrical and movement artifacts. This stage was recorded for 27 h including the first 3 h of habituation and 24 h of recording, from 13:30 to 16:30. Signals from the LPa and EMG electrode, referenced to the cerebellum electrodes, were amplified and filtered (2 000 and 5 000 gain, 0.1 and 0.003 s time constant for EEG and EMG, respectively; high cut-off filter for both; AB-621G, Nihon-Koden, Tokyo, Japan) and digitized at a 500 Hz sampling rate (Power 1401; Cambridge Electronic Design, Cambridge, UK).

### Sleep scoring

Sleep/wake stages were scored using the automatic scoring tool “rat sleep auto” script in Spike2 (Cambridge Electronic Design). Scores were calculated by analyzing the EEG and EMG signals in consecutive 30 s epochs. We also manually confirmed the scoring procedure accuracy using EEG derivatives and EMG signals to segregate the WAKE (desynchronized EEG and medium-high amplitude EMG), NREM (high-voltage slow-wave EEG and low-amplitude EMG), and REM (EEG theta activity and very low amplitude EMG) stages for each epoch during the 24 h recording period [3].

### Statistical analyses

Statistical analyses were mostly performed using GraphPad Prism 8 (La Jolla, CA, USA). For statistical comparisons of two groups, the Mann–Whitney U test, as the nonparametric version of the parametric t-test, was used. For comparisons of more than two groups, analysis of variance (ANOVA) was used followed by post hoc tests. Alpha diversity was calculated using three different parameters: Chao1 richness estimator and Shannon and inverse Simpson indexes. Principal coordinate analysis was performed using XLSTAT software package (Addinsoft, Long Island City, NY, USA). All data represent the mean ± SEM.

## Results

### Gut microbiota analysis

We studied fecal gut microbiota before (before), during (stress 5d, 11d), and after (after stress, 1W, 1M) the 14 day SDS. SDS significantly increased the levels of fecal classes Betaproteobacteria and Flavobacteriia, while decreasing those of Clostridia (Fig. 1A, after stress). Bacteroidia and Bacilli showed a tendency to increase, whereas *Actinobacteria* tended to decrease.

**Fig. 1.**
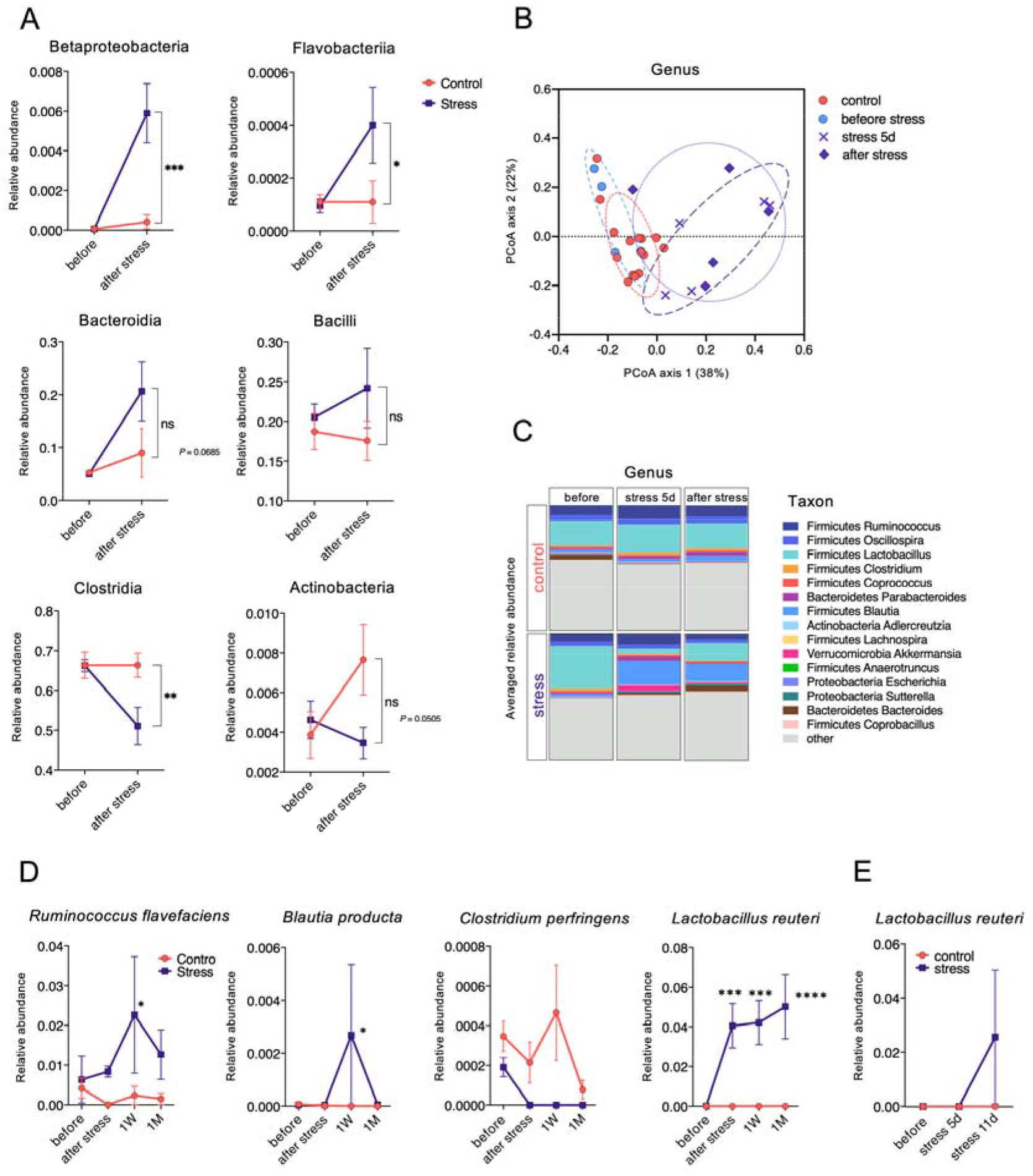
Changes in the fecal microbiome by SDS. **A** Class-level alterations in fecal samples of control and SDS rats before and one day after SDS. \**P* < 0.05, \*\**P* < 0.005, \*\*\**P* < 0.0005, two-way (day × group) ANOVA followed by Sidak’s multiple comparisons test. **B** Genus-level principal coordinate analysis (PCoA) plot based on Bray–Curtis distances. The plot demonstrates a significant shift in control, pre-stress baseline (before stress), day 5 (stress 5d), and SDS (after stress) groups. **C** Relative abundance at the genus level before, during (stress 5d), and after SDS. **D** Relative abundance at the species level before (before) and 1 day (after stress), 1 week (1W), and 1 month (1M) after SDS. \**P* < 0.05, \*\*\**P* < 0.0005, \*\*\*\**P* < 0.0001, two-way (day × group) ANOVA followed by Sidak’s multiple comparisons test. **E** Relative abundance of *L. reuteri* before (before) and during (stress 5d, 11d) SDS. ns, not significant.

Alpha diversity was analyzed to assess differences in within-subject diversity. The Chao1 richness estimator and Shannon and inverse Simpson indexes showed no statistically significant changes during and after SDS (Supplementary Fig. 1A-C). Analysis of beta diversity at the genus level was conducted to determine how the whole microbial community changed chronologically. Beta diversity exhibited no significant difference as assessed by Bray–Curtis distances before SDS (Fig. 1B; before). Bray–Curtis distances increased in the SDS rats at day 5 (Fig. 1B; stress 5d). Following SDS completion (Fig. 1B; after stress), Bray–Curtis distances further increased significantly (*P* < 0.0001).

Genus-level microbiota structure also markedly changed during the SDS period (Fig. 1C). When compared to before stress (before), *Lactobacillus* showed clear decreases at day 5, whereas *Blautia* exhibited large increases. These trends were fairly consistent after the last SDS exposure.

At the species level, *L. reuteri* levels significantly increased following SDS (Fig. 1D), with further increases observed even one month after SDS completion (Fig. 1D). Conversely, other species (*Ruminococcus flavefaciens*, *Blautia producta*, and *Clostridium perfringens*) exhibited only temporary change (Fig. 1D). More precise chronological analysis during the SDS period further revealed an increase in *L. reuteri* levels later in the SDS period (stress 11d) (Fig. 1E).

### Sulfur metabolite analyses

Next, we analyzed SDS-induced changes in the fecal sulfur metabolites. The Sulfur index before and after SDS revealed significant decreases of glucose, methionine, and serine levels in the SDS rats (Fig. 2A). However, only ergothioneine, a natural thiol compound present in mushrooms and mammalian tissues [21, 22], exhibited a significant increase following SDS. We then evaluated chronological changes in the fecal ergothioneine levels in the SDS rats. A substantial increase in the amount of fecal ergothioneine was observed at day 11 (Fig. 2B). Statistically significant increases in the fecal ergothioneine levels were maintained even one month after the last SDS (Fig. 2B).

**Fig. 2.**
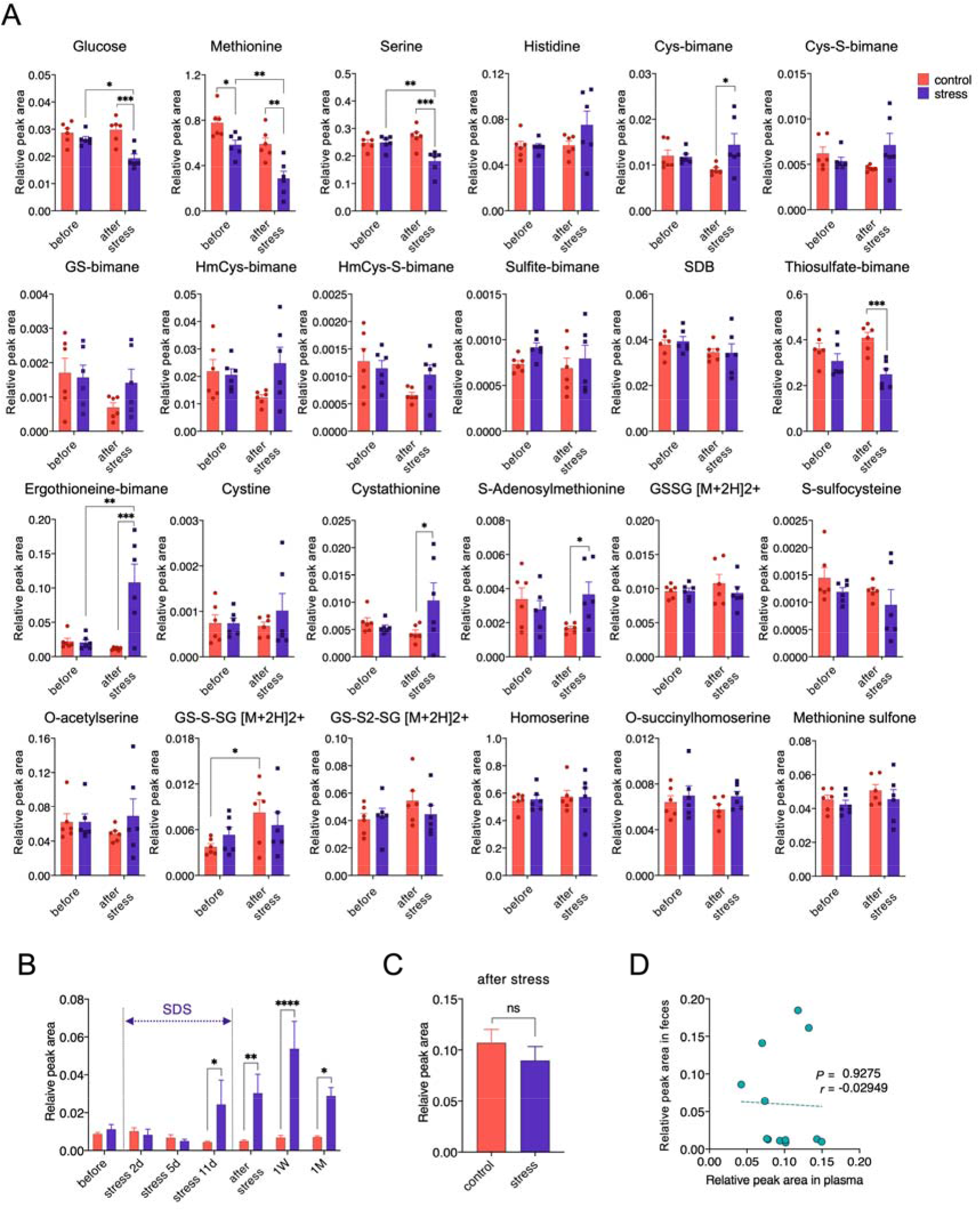
Changes in the fecal and plasma sulfur metabolites by SDS. **A** Changes in the levels of fecal metabolites before and one day after SDS. In total, 24 sulfur metabolites could be measured in the rat fecal samples. \**P* < 0.05, \*\**P* < 0.005, \*\*\**P* < 0.0005, two-way (day × group) ANOVA followed by Sidak’s multiple comparisons test. Cys, cysteine; GS, S-conjugate of glutathione; HmCys, homocysteine; SDB, sulfide with monochlorobimane; GSSG, glutathione disulfide; GS-S-SG, a persulfide form of glutathione disulfide. **B** Time course of fecal ergothioneine levels. Fecal ergothioneine relative peak area was measured before, during (stress 2d, 5d, 11d), and after SDS (1 day: after stress, 1 week: 1W, 1 month: 1M). *F* (6, 42) = 6.849, *P* < 0.0001, \**P* < 0.05, \*\**P* < 0.005, \*\*\*\**P* < 0.0001, two-way (day × group) ANOVA followed by Sidak’s multiple comparisons test. **C** No significant difference was observed in the plasma ergothioneine levels (relative peak areas) between control and SDS rats one day after SDS. *P* = 0.3095, Mann–Whitney U test. n = 6/group. **D** No significant correlation existed between plasma and fecal ergothioneine levels one day after SDS. ns, not significant.

We also evaluated whether the fecal ergothioneine levels coordinated with plasma levels. Plasma ergothioneine levels one day following SDS did not significantly differ between control and SDS rats (Fig. 2C). Furthermore, plasma and fecal ergothioneine levels did not significantly correlate (Fig. 2D), indicating that although SDS increases fecal ergothioneine levels, ergothioneine may not be absorbed by the large intestinal tract, or the fecal levels may be insufficient to increase plasma levels.

### Production of ergothioneine by *L. reuteri*

As ergothioneine biosynthesis occurs only in fungi or mycobacteria but not in mammals [23, 24], and chronological changes in *L. reuteri* and ergothioneine fecal levels were comparable in SDS rats, we hypothesized that the increase in the fecal ergothioneine levels in the SDS rats may be related to changes in gut microbiota. We therefore comprehensively examined the correlation between all gut bacterial species and all evaluated sulfur metabolites one day following SDS. Among gut bacteria, *L. reuteri* levels most significantly correlated with ergothioneine (Fig. 3A; *P* < 0.0001, *r* = 0.82). Moreover, the relative abundance of *L. reuteri* significantly positively correlated with fecal ergothioneine amounts when all chronological samples were analyzed together (Fig. 3B; *P* < 0.0001, *r* = 0.8310).

**Fig. 3.**
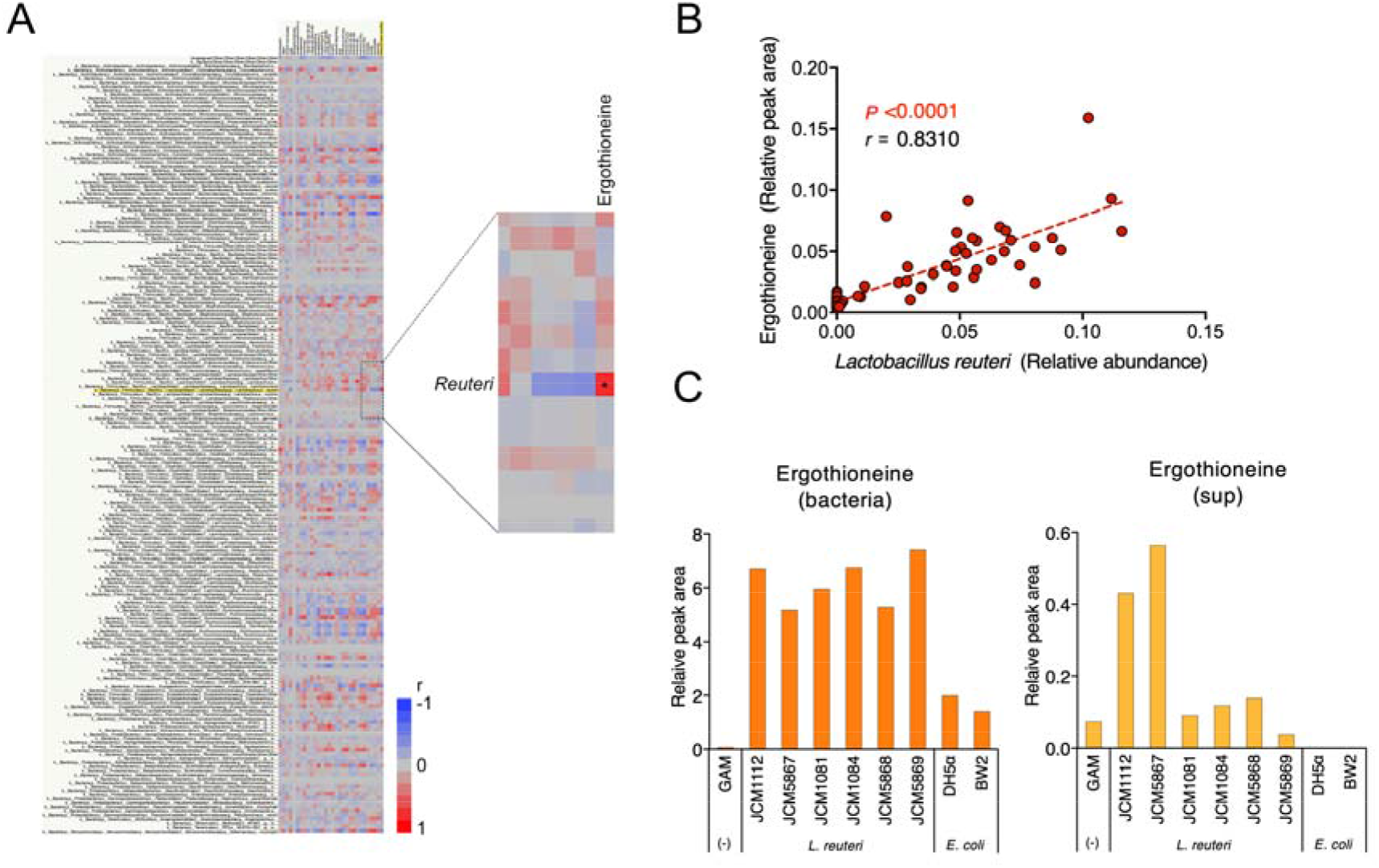
Ergothioneine production by *L. reuteri*. **A** Correlation analysis of fecal gut bacterial (species level) and sulfur metabolites. The enlarged image shows a strong positive correlation in the levels between the gut bacterium *L. reuteri* and the sulfur metabolite ergothioneine. **B** Significant positive correlation between fecal *L. reuteri* bacterial relative areas and ergothioneine amounts. *P* < 0.0001, *r* = 0.8310. **C** Ability of *L. reuteri* to produce ergothioneine in vitro. Relative peak areas of ergothioneine in the bacterial bodies (left; bacteria) and supernatants (right; sup) of *L. reuteri* or *E. coli* are shown.

To directly demonstrate that *L. reuteri* produces ergothioneine, we conducted *L. reuteri* and *E. coli* culture experiments. Ergothioneine levels in the bodies and supernatants of *L. reuteri* were much higher than those of *E. coli* (Fig. 3C).

### Effects of L-ergothioneine administration on the SDS-induced depressive behavior and sleep abnormalities

Ergothioneine has been shown to accumulate at high concentrations (100 µM to 2 mM) in most cells and tissues of mammals including the brain [25–27]. Carnitine/organic cation transporter 1 (OCTN1) [28], encoded by the gene *SLC22A4*, plays a pivotal role in this intracellular accumulation. Indeed, silencing this gene completely inhibits uptake of ergothioneine [27, 29]. As the ileum expresses OCTN1 most abundantly in the body [28, 30], dietary ergothioneine is considered to be mainly absorbed in the ileum [22, 27] and is subsequently transported across the blood-brain barrier into the brain [31]. We therefore evaluated whether the protective properties of ergothioneine extend to the whole organism upon oral administration in SDS rats. We orally administered 0.25 mg/ml L-ergothioneine as a preventative measure 7 days prior to SDS (day −7) and continued oral ingestion until the end of the SDS period (Fig. 4A). Interaction tests immediately following SDS showed that SDS rats exhibited significant social avoidance behaviors against unfamiliar BN or SD rats (Fig. 4B). Preventative L-ergothioneine administration resulted in statistically significant improvements in the avoidance behaviors against resident BN rats (*P* = 0.0198) and almost significant improvements against SD rats (*P* = 0.0542) immediately after SDS (Fig. 4B; after stress). These significant improvements in avoidance behaviors against BN rats were still observed one month after the last SDS (*P* = 0.0130, Fig. 4B; 1M).

**Fig. 4.**
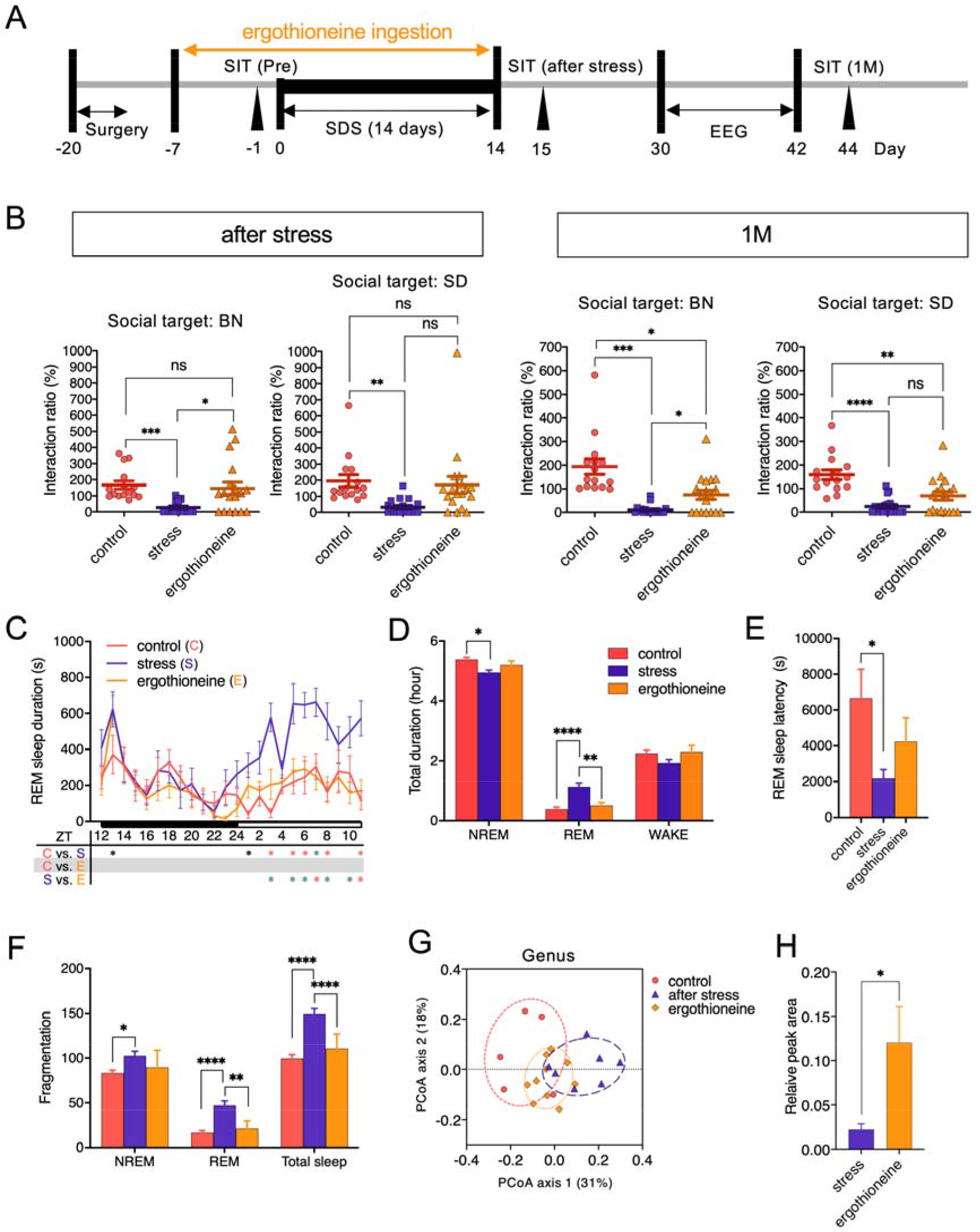
Antidepressant effects of oral L-ergothioneine ingestion in the SDS rat model. **A** Experimental design of oral L-ergothioneine ingestion, SDS session, social interaction test, and 24 h EEG recording. **B** Social interaction test one day or one month following SDS (day 15 or day 44 in Fig. 4A, n = 15–18/group). \**P* < 0.05, \*\**P* < 0.005, \*\*\**P* < 0.0005, \*\*\*\**P* < 0.0001, one-way ANOVA (Welch and Browne–Forsythe tests) followed by Dunnett’s T3 multiple comparisons test. **C** REM sleep variation. *F* (46, 506) = 2.260, *P* < 0.0001, \**P* < 0.05 (black), \**P* < 0.005 (green), \**P* < 0.001 (red), two-way (time × group) repeated ANOVA followed by Tukey’s multiple comparisons test. ZT, Zeitgeber time. n = 7–10/group. **D** Total duration in NREM, REM sleep, and wakefulness (WAKE) during the light phase [08:00 (ZT0) – 16:30 (ZT8.5)]. *F* (4, 66) = 9.613, *P* < 0.0001, \**P* < 0.05, \*\**P* < 0.005, \*\*\*\**P* < 0.0001, two-way (sleep stage × group) ANOVA followed by Tukey’s test. n = 7–10/group. **E** REM sleep latency. *F* (2, 22) = 4.033, *P* = 0.0322, \**P* < 0.05, one-way ANOVA followed by Tukey’s test. **F** Total number of fragmentations of NREM, REM sleep, and WAKE during the light phase (ZT0–ZT8.5). *F* (4, 66) = 3.000, *P* = 0.0245, \**P* < 0.05, \*\**P* < 0.005, \*\*\*\**P* < 0.0001, two-way (sleep stage × group) ANOVA followed by Tukey’s test. n = 7–10/group. **G** Recovery of the abnormal shift in the SDS rat microbiota by L-ergothioneine. **H** Fecal ergothioneine levels following L-ergothioneine oral administration. *P* = 0.0185, Mann–Whitney U test.

SDS also induced significant abnormalities in sleep such as increased REM sleep duration (Fig. 4C, D), shortened REM latency (Fig. 4E), and increased numbers of fragmentations (Fig. 4F) during the light phase. L-ergothioneine ingestion significantly improved these REM sleep abnormalities except for the REM sleep latency. Moreover, in NREM sleep, SDS significantly decreased the duration (Fig. 4D) and increased the fragmentation number (Fig. 4F), whereas L-ergothioneine tended to improve these abnormalities.

Finally, microbial diversity was analyzed in the feces from control, stressed, and L-ergothioneine-treated rats. Alpha diversity in the feces of SDS rats tended to be lower (*P* = 0.0582) based on the Chao1 estimator when compared to that of controls, but other indexes (Shannon and inverse Simpson) showed no statistically significant difference among the three groups (Supplementary Fig. 1D-F).

Beta diversity analysis at the genus level revealed that L-ergothioneine administration has a preventative effect on the abnormal shift of the whole microbial community induced by SDS (Fig. 4G). L-ergothioneine administration significantly increased the levels of fecal ergothioneine (approximately five times) (Fig. 4H), suggesting that biologically relevant amounts of ergothioneine were most likely absorbed from the ileum via OCTN1 in our paradigm, and the remainder was excreted in the feces.

## Discussion

The major findings in this study were as follows. 1) Increases in the amount of fecal *L. reuteri* were observed 11 days after the initiation of SDS. The levels of *L. reuteri* continued to increase significantly and were retained at least for one month following the last SDS. 2) The Sulfur index showed significant increases in the levels of fecal ergothioneine 11 days after SDS initiation. These increases peaked at one week and remained significantly increased one month after the last SDS. 3) The increased amounts of fecal *L. reuteri* significantly positively correlated with the ergothioneine levels. We further found that *L. reuteri* directly produced ergothioneine in vitro. 4) Given the anti-inflammatory and anti-oxidative actions of ergothioneine, we orally administered L-ergothioneine to the SDS rats as a preventative measure and found that the social avoidance behaviors were ameliorated. We additionally observed improvements in sleep abnormalities, particularly those relating to REM sleep.

Previous studies showed that the feces transplanted from patients with MDD caused MDD-like behavior in germ-free animals [10, 11], suggesting that the proportion of harmful “depression bacteria” may predominate in the feces of these patients [11]. At the genus level, *Coprococcus* and *Dialister*, so-called “depression bacteria”, are known to increase in the feces of patients with MDD and induce MDD-like behaviors in animals [32]; however, we found no such changes related to these bacteria in our study (Fig. 1C). Rather, we found a quite unique long-lasting increase in the levels of *L. reuteri* in our SDS rat model (Fig. 1D, E). *L. reuteri*, a potential probiotic known to modulate the immune system [33], may decrease anxiety as measured on the elevated plus maze [34], reduce the stress-induced increase of corticosterone levels [35], and reduce despair-like behavior in mice [36, 37]. A similar increase in the level of *L. reuteri* was noted in a rat SDS model with a 7day SDS regimen [13]. Resilient rats showed a greater increase in *L. reuteri* levels than did vulnerable rats; however, further mechanistic studies were not performed [13].

In the present study, we also found a significant sharp increase in the levels of the sulfur metabolite ergothioneine in the feces of SDS rats in the second half of the SDS period, with this characteristic increase also continuing for at least a month following the cessation of stresses. The biosynthesis of ergothioneine, a highly stable anti-oxidative and anti-inflammatory sulfur metabolite [23, 38–41], occurs only in fungi or mycobacteria [23, 24], suggesting that the changes in the gut microbiota may cause such increase. As expected, comprehensive correlation analysis demonstrated that fecal ergothioneine and *L. reuteri* levels significantly correlated. In addition, in vitro study further demonstrated that *L. reuteri* exhibits considerably greater ergothioneine production capability than does *E. coli*. Given the well-documented anti-inflammatory effects of *Lactobacillus* [33] and ergothioneine [23, 38–41], these findings suggest that an anti-inflammatory defense mechanism mediated via *L. reuteri* and its sulfur metabolite ergothioneine may occur in the lower gastrointestinal tract as a result of long-term psychosocial stress. This concept was further supported by preliminary experiments involving fecal S100A9 (Supplementary Fig. 2), a protease-resistant neutrophil-derived protein [42–44] that can be used to predict relapse of inflammatory bowel diseases [44–46]. We found that the levels or rate of increase of S100A9 was elevated in the feces of SDS rats but returned to control levels by oral ergothioneine administration (Supplementary Fig. 2), suggesting the occurrence of bowel inflammation due to SDS and anti-inflammatory effects of ergothioneine. In our rat model, however, this defense mechanism may not be efficiently reflected in the body as no difference was observed in the plasma ergothioneine levels.

In addition to its anti-oxidative actions, ergothioneine also exerts several cytoprotective functions such as lipid peroxidation inhibition [47, 48] and DNA/protein damage control [23, 49]. A study using control mice reported the antidepressant-like effects of ergothioneine in forced swim and tail suspension tests [31]; however, to our knowledge, no study has examined the effect of ergothioneine in stressed animal models. We therefore investigated the effects of oral ergothioneine administration on MDD-like behavior and sleep abnormalities in our SDS rat model and confirmed its preventative effects. Although the functional mechanism associated with the antidepressant effect of ergothioneine remains unknown, mounting evidence suggests that increased central or peripheral inflammatory processes may be involved in stress-related mood disorders, such as MDD [31, 50–53]. Consistent with this, inflammatory cytokines are elevated in patients with depression [54–57]. Furthermore, numerous animal studies have shown that inflammatory mechanisms underlie stress-induced depressive-type behaviors [54–57]. Therefore, it is plausible that, following absorption, ergothioneine may manifest its antidepressant effects through anti-inflammatory mechanisms of the central and/or peripheral nervous system.

Several possibilities underlying ergothioneine-mediated rescue of sleep abnormalities should be mentioned. First, neuronal circuit-level dysregulation of the sleep stage transition in the brainstem region [58–60] occurred consequent to SDS-associated inflammation, which may be prevented by the anti-inflammatory effects of ergothioneine as described above. Second, abnormalities in endogenous sleep substances (e.g., prostaglandin D2 or adenosine), including reduced glutathione, which plays an important role in removing reactive oxygen species, may also be involved in the sleep abnormalities [61, 62]. For example, reduced amounts of glutathione were observed in the brains of acutely sleep-deprived rats [63, 64]. More recently, a study using short-sleeping *Drosophila* mutants revealed a bidirectional relationship between oxidative stress and sleep: oxidative stress triggers sleep, which then acts as an antioxidant for both the body and the brain [65]. In this context, the concept of treatment and prevention of depression-related sleep abnormalities by controlling intracerebral oxidative stress production may be ground-breaking, as ergothioneine may harbor enormous potential as a direct sleep improvement tool.

Limitations of this study should be noted. First, the mechanisms underlying improvement of the MDD-like symptoms in the SDS rats by oral ergothioneine administration remain unelucidated. It is plausible that oral ergothioneine simply ameliorates SDS-induced inflammatory events in the periphery (especially in the intestinal flora), which in turn inhibits the MDD-like symptoms via unknown mechanisms. Second, we did not produce any CNS data related to disturbed sleep and inflammation. This is partly because of the lack of knowledge regarding the CNS center responsible for disturbed sleep associated with MDD, a major behavioral phenotype recapitulated in the current model. Nonetheless, we focused on the septohippocampal pathways that generate the hippocampal REM theta oscillation [66], since REM theta powers most significantly correlated with the MDD-like social interactions in our SDS rats treated with antidepressants [3] and L-ergothioneine (Supplementary Fig. 3). Further study is warranted to clarify the related CNS changes. Finally, we cannot exclude the possibility that the bacterial flora of SDS-administered SD rats was influenced by that of BN rats, considering that they had direct contact.

In conclusion, for the first time, we demonstrated the role of *L. reuteri* and identified its sulfur metabolite ergothioneine as a candidate molecular inhibitor of psychosocial stress in the microbiota-gut-brain axis [4–9]. Oral L-ergothioneine ingestion significantly prevented MDD-like social avoidance and sleep abnormalities in our SDS rats. Considering that ergothioneine is a natural ingredient that is incorporated into the blood via OCTN1 mainly in the ileum [20], our data warrant future evaluation of preventative use of ergothioneine in stress-related diseases including MDD. Furthermore, increases in fecal ergothioneine may serve as an effective MDD biomarker, at least for a subtype.

## Acknowledgments

The authors thank Ms. Moe Watanabe for her technical assistance. We would like to thank Norihito Murayama and Hiroshi Watanabe from the Suntory Global Innovation Center for their technical suggestions. This study was funded by JSPS KAKENHI Grant Numbers 25871162 and 16K04442 to YM and 15K15438 to YT.

## Author contributions

YM, KW, KS, and YT designed the experiments; YM, NO, TS, KW, YK, IO, and YT performed the experiments; YM, KW, KS, and YT designed the analyses and discussed the results; YM, KW, KS, YK, IO, and YT analyzed the data; YM and YT wrote the paper; YM and NO performed animal surgeries; and all authors commented on the manuscript.

