## supplementray information for "Ergothioneine, a metabolite of the gut bacterium *Lactobacillus reuteri*, protects against stress-induced sleep disturbances"

**This file includes:**

Supplementary Figure 1

Supplementary Figure 2

Supplementary Figure 3

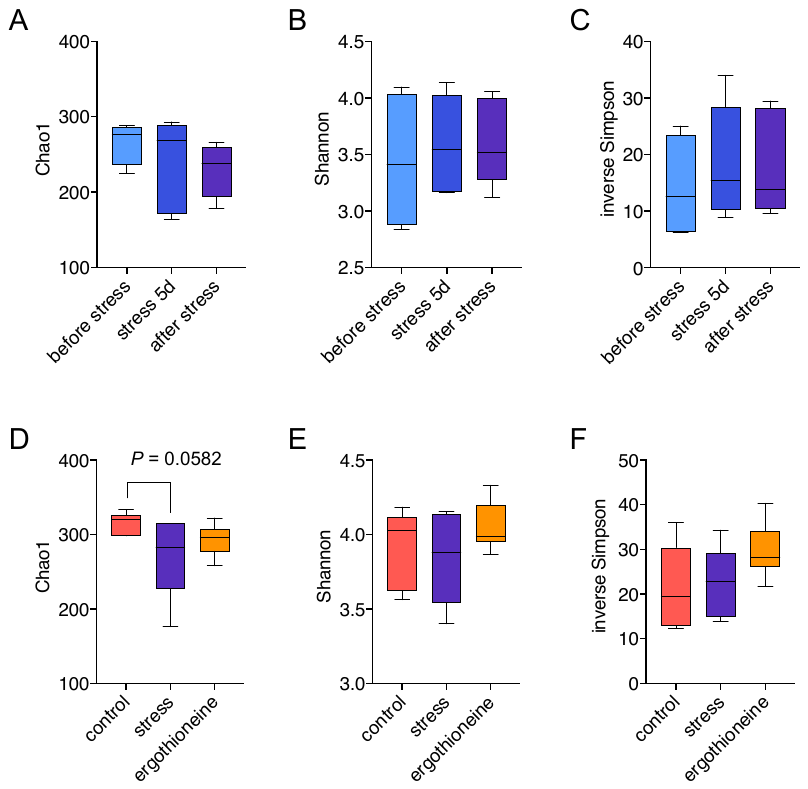

**Supplementary Figure 1	Alpha diversity of fecal microbiota.**

A-C. Chao1 richness estimator (A) and Shannon (B) and inverse Simpson (C) indexes showed no significant differences during and after stress. Data are expressed as the median. Whiskers show the minimum and maximum values.
D-F. Chao1 richness estimator (D) tended to be lower (*P* = 0.0582, ordinary one-way ANOVA followed by Tukey’s test) in microbiota of SDS rats relative to control rats. The other indicators of alpha diversity [Shannon (E) and inverse Simpson (F) index] showed no significant differences among the three groups.

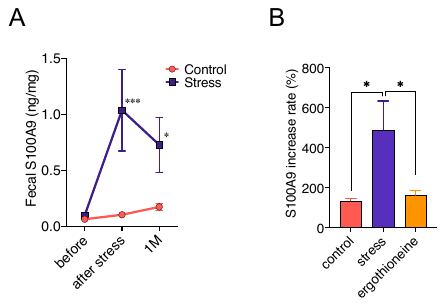

**Supplementary Figure 2	Increased fecal S100A9 levels in SDS rats.**

A. Effects of SDS on fecal S100A9 levels. Calprotectin, a heterodimer of S100A8 and S100A9, has been reported to be a useful biomarker in inflammatory bowel disease [53, 54]. We measured fecal S100A9 levels before and one day (after stress) and 1 month (1M) after the last SDS using an enzyme-linked immunosorbent assay (ELISA) for S100A9 (Yamasa Corp., Chiba, Japan) [55]. Fecal S100A9 levels significantly increased 1 day (after stress) and even 1 month (1M) after the last SDS, suggesting that SDS induces prolonged inflammation in the intestinal tract of rats. **P* < 0.05, ****P* < 0.001.
B. Effects of ergothioneine on the rate of increase of fecal S100A9 levels in SDS rats. Preventative administration of ergothioneine restored the rate of increase of fecal S100A9 levels. Data are shown as the rate of increase (%) of fecal S100A9 levels before and after SDS (after/before x 100). **P* < 0.05.

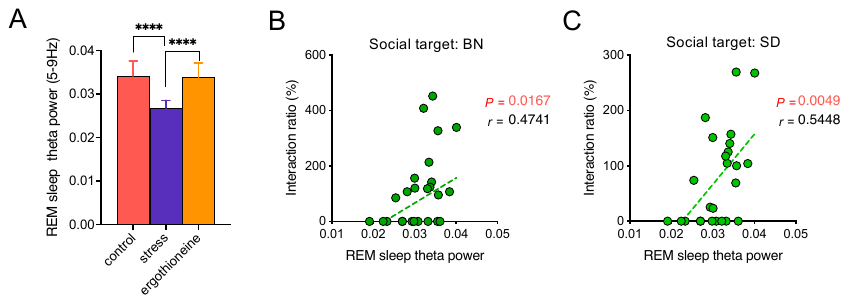

**Supplementary Figure 3 Increased theta activity by oral ingestion of L-ergothioneine.**

A. Averaged relative spectral power of theta (5–9 Hz) during REM sleep of the light phase (ZT0 – ZT8.5). EEG power spectrum analysis was performed to investigate the effects of oral L-ergothioneine ingestion on theta activity, which reflects pyramidal neuronal activity in dorsal and posterior dorsal CA1. Individual relative power spectra per one epoch for REM sleep of the light phase were calculated as the ratio of each power spectrum value to the sum of the powers of the band and then averaged [3]. SDS significantly reduced EEG theta power during REM sleep, but oral ingestion of L-ergothioneine significantly restored the reduced theta power to control levels, suggesting that L-ergothioneine treatment remarkably enhances the generation of theta waves in the brain of SDS rats. *F* (30, 352) = 2.685, *****P* < 0.0001, two-way (group x frequency) ANOVA followed by Tukey's multiple comparisons test.
B-C. Correlations between averaged relative REM theta power during the light phase (ZT0–ZT8.5) and interaction ratios corresponding to BN rat (B) or SD rat (C) social targets one month after the last SDS. REM theta power significantly correlated with social interaction ratios (BN; *P* = 0.0167, SD; *P* = 0.0049).
